# Mesenchymal stem cells carry and transport clusters of cancer cells

**DOI:** 10.1101/2021.02.11.430875

**Authors:** Jana Zarubova, Mohammad Mahdi Hasani-Sadrabadi, Sam CP Norris, Andrea M Kasko, Song Li

**Affiliations:** Department of Bioengineering, University of California, Los Angeles, CA 90095, USA; Department of Biomaterials and Tissue Engineering, Institute of Physiology of the Czech Academy of Sciences, Prague, 14220, Czech Republic

## Abstract

Cell clusters that collectively migrate from primary tumors appear to be far more potent in forming distant metastases than single cancer cells. A better understanding of collective cell migration phenomenon and the involvement of different cell types during this process is needed. Here, we utilize a micropatterned surface composed of a thousand of low-adhesive microwells to screen motility of spheroids containing different cell types by analyzing their ability to move from the bottom to the top of the microwells. Mesenchymal stem cells (MSCs) spheroid migration was efficient in contrast to cancer cell only spheroids. In spheroids with both cell types mixed together, MSCs were able to carry the low-motile cancer cells during migration. As the percentage of MSCs increased in the spheroids, more migrating spheroids were detected. Extracellular vesicles secreted by MSCs also contributed to the pro-migratory effect exerted by MSCs. However, the transport of cancer cells was more efficient when MSCs were physically present in the cluster. Similar results were obtained when cell clusters were encapsulated within a micropatterned hydrogel, where collective migration was guided by micropatterned matrix stiffness. These results suggest that stromal cells facilitate the migration of cancer cell clusters, which is contrary to the general belief that malignant cells metastasize independently.

**Significance:** During metastasis, tumor cells may migrate as a cluster, which exhibit higher metastatic capacity compared to single cells. However, whether and how non-cancer cells contained in tumor cluster regulate it’s migration is not clear. Here, we utilize two unique approaches to study collective tumor cell migration in vitro: first, in low-adhesive microwells and second, in micropatterned hydrogels to analyze migration in 3D microenvironment. Our results indicate that MSCs in tumor cell clusters could play an important role in the dissemination of cancer cells by actively transporting low-motile cancer cells. In addition, MSC-released paracrine factors also increase the motility of tumor cells. These findings reveal a new mechanism of cancer cell migration and may lead to new approaches to suppress metastases.

## Introduction

The majority of cancer-related deaths are not caused by the primary tumor, but rather by the metastatic spreading of cancer cells to distant organs (1). Metastasis is generally described as a multistep process, in which transformed epithelial cells acquire mesenchymal features, detach from the primary tumor, migrate through a tissue to the vasculature and travel to distant sites where they proliferate (2). Until recently, metastases were thought to originate from single cancer cells, however, recent research suggests that up to 97% of metastases arise from clusters of cells that collectively migrate from the primary tumor, circulate in the blood and extravasate at distant sites (3–6). These circulating tumor cell clusters (CTCCs) contain between 4 – 100 cells (7) and may also contain mesenchymal cells, endothelial cells, and/or immune cells (8).

Collective cell migration may facilitate metastasis for several reasons. Cells are better protected from shear forces within the vasculature and an immune cell attack. Although CTCCs are larger than single cells, they can still dynamically reorganize and adjust their geometries to pass through narrow blood vessels (7). Co-traveling cells may also create a favorable environment that supports cancer cell survival and proliferation at the early phases of metastatic growth (9). The importance of co-traveling stromal cells in the metastatic process is supported by observations that up to 86% of carcinoma cells that spread to the lungs were accompanied by primary tumor stroma-derived cells and that these cells, in majority of cases, stained positive for smooth muscle α- actin (αSMA) but only in 28% cases for F4/80, a macrophage marker (9). When the tumor stromal cells were partially depleted, the number of metastases significantly decreased. Thus, cancer progression does not depend solely on cancer cells but also on the tumor microenvironment that can change dramatically during the evolution of the disease.

While mesenchymal stem cells (MSCs) are known to be recruited to damaged or inflamed tissue, where they promote tissue regeneration, MSCs are also attracted to tumor sites (10–12). This homing ability has been utilized where MSCs act as delivery vehicles for anti-cancer therapies (13). However, the use of MSCs for tumor targeting or their application in regenerative therapies after tumor resection can be potentially detrimental. MSCs and the extracellular vesicles (14, 15) they secret may increase cancer growth in various ways, such as: by promoting angiogenesis (16, 17); by suppressing immune responses through induction of regulatory T cells (18); or by polarizing macrophages towards pro-tumorigenic M2 phenotype (19). Moreover, there is an evidence that MSCs can differentiate into cancer-associated fibroblasts (CAFs) (20) that might increase cancer invasiveness by secreting matrix metalloproteinases (MMPs) (21), which are responsible for the degradation of extracellular matrix (ECM) components and release of growth factors bound to the ECM. Studies also show that MSCs can dramatically increase cancer metastatic potential when they are in close proximity to cancer cells. This is mediated by the release of soluble factors such as chemokine CCL5 (22, 23) or growth factors, such as TGF-β (24) that promote the epithelial-to-mesenchymal transition (EMT) and increase cancer cell motility. However, it is not clear what role MSCs (or stromal cells, in general) have in the CTCCs. It is still unknown if MSCs contribute to the increased metastatic incidence mainly by production of paracrine factors that promote cancer EMT and provide a favorable niche at the secondary metastatic organs or if there are other ways by which these cells participate in the metastatic dissemination of CTCCs.

To understand the role of MSCs in the collective migration of CTCCs, we develop an *in vitro* model that enables us to follow and quantify the migratory ability of hundreds of cell clusters at once. Our results suggest that the current theory of malignant cells bringing their own soil (passenger stromal cells) to the secondary metastatic sites might not be accurate. Instead, stromal cells may actively facilitate cancer cell migration and metastasis.

## Results

### Collective migration of cell clusters is cell-type dependent

To analyze differences in the collective migration of cell clusters formed by different cell types *in vitro*, cells were seeded as single-cell suspension into low-adhesive inverted-pyramidal microwell plates at a concentration of 50 cells per microwell to form aggregates. This technique has been used to fabricate cellular aggregates based on various cell types for *in vitro* cellular studies (25), fabrication of therapeutics (26), and drug screening (27). While forming cell aggregates, we observed that some aggregates migrated out of the microwells. When the surface was incubated with an anti-adhesive solution for a prolonged period of time, a completely anti-adhesive surface was created that didn’t promote any migration. However, short incubation times (10min) with an anti-adhesive solution enabled the formation of a low-adhesion surface that promoted formation of spheroids but also permitted cell cluster migration. If no anti-adhesive coating was applied, cells spread on the surface and did not form spheroids. If pre-formed spheroids were seeded on the non-coated microwells, they spread and did not move. To test if the spheroid migration behavior in the microwell plates was unique to the microwell topology, we examined the motility of pre-formed fibroblast spheroids on low adhesive flat surfaces. Within micropatterned surfaces, the migrated distance and velocity of cell aggregate migration was significantly greater than the flat surface (**Fig. 1A**). Therefore, we used the micropatterned surfaces for the remainder of this study.

**Fig. 1.**
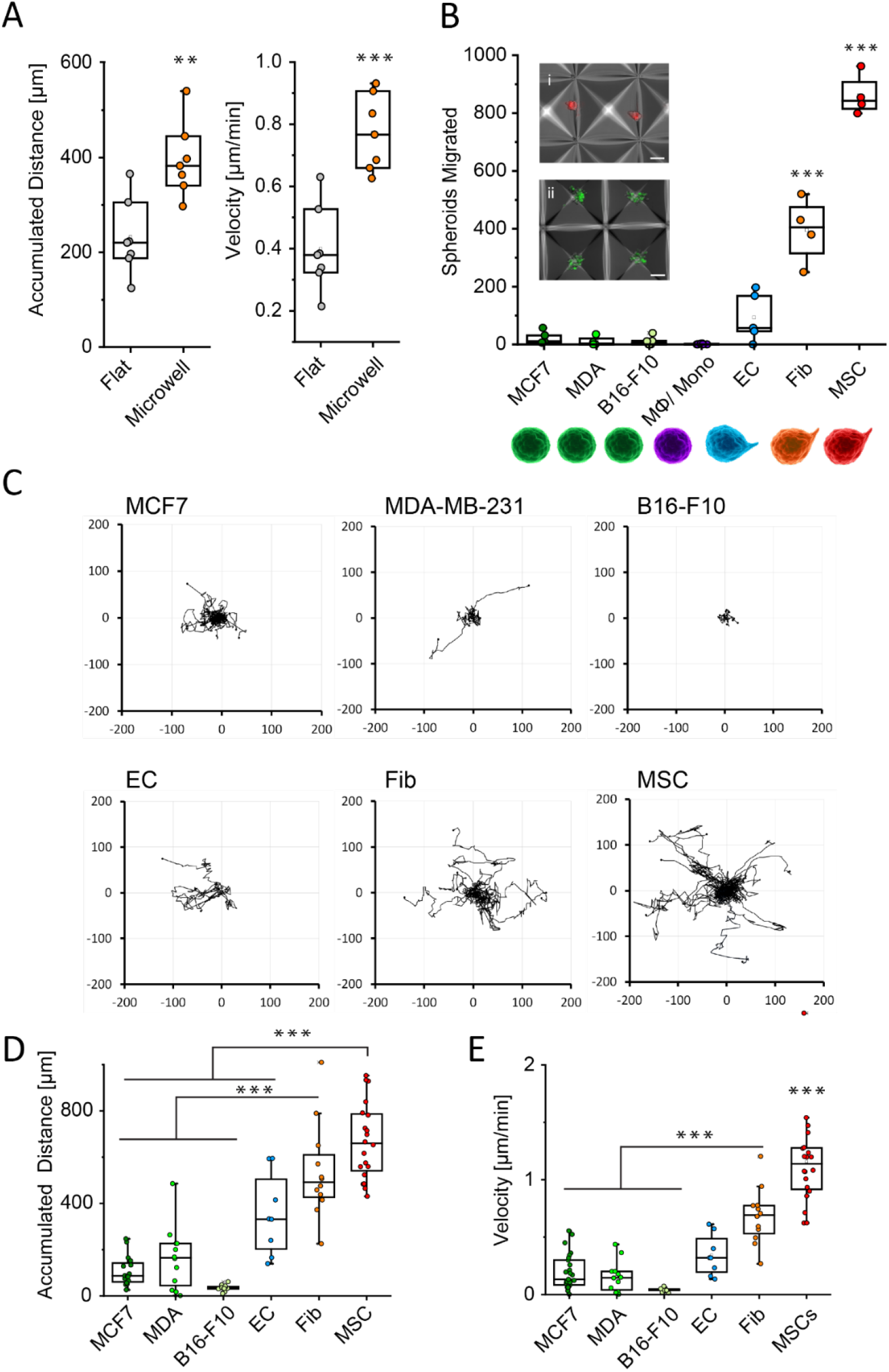
Differences in the migration of spheroids formed by single cell types. **A.** Accumulated distance and velocity of fibroblast spheroids migrating on flat low-adhesive surface and in the micropatterned surface. **B**. Number of spheroids migrated to the top of low-adhesive microwells after 24 hours for breast cancer cell lines (MCF7 and MDA-MB-231), melanoma cancer cell line B16-F10, monocytes or macrophages, endothelial cells (EC), fibroblasts (Fib), MSCs. i - MSC spheroids migrated to the top of the microwells after 24 hours; ii - cancer cell aggregated in the low-adhesive microwells after 24 hours with no displacement observed, scale bar - 100 μm. **C**. Migration trajectories of spheroids in the microwells evaluated from time-lapse microscopy videos. **D**. Total distance migrated by the spheroids. E – comparison of speed of different spheroids formed by single cell types. *** p < 0.001, ** p < 0.01, * p < 0.05.

Depending on the cell composition of the aggregates, differences in cluster migration from the bottom to the top edge of the microwells were observed. Displacement of the spheroids was evaluated after 24 hours. Each well of a 24 well plate contained approximately 1000 microwells, which allowed for the evaluation of 1000 cell clusters simultaneously. In this study, migration of spheroids formed by three cancer cell lines (MCF7, MDA-MB-231, and B16-F10), macrophages, endothelial cells (ECs), fibroblasts and MSCs was tested. Of these cells, MSC spheroids showed the highest migratory ability. The majority of MSC spheroids reached the top of the microwell after 24 hours (**Fig. 1B**). Interestingly, all displaced spheroids remained at the top edges of the microwells (**Fig. 1Bii**) and did not migrate back to the bottom.

**Fig. 1C** shows analyses of spheroid motility in the low-adhesive microwells evaluated from time-lapse microscopy videos (**Supplemental videos S1 and S2**). Pyramidal microwells approximately 400 μm in length, 400 μm in width and 300 μm in depth were used for a majority of the experiments performed (**Supplemental Fig. 1**). Larger-sized microwells were used to test the migratory behavior of bigger spheroids (*e.g.*, 2000 cells), which resulted in migration comparable to the smaller cell clusters (**Supplemental video S3**). We did not observe any significant differences in the spheroid migration between the low metastatic breast cancer cell line (MCF7) and highly metastatic breast cancer cell lines (MDA-MB-231) and melanoma (B16-F10). Likewise, EC clusters moved significantly less than fibroblast and MSC clusters, which migrated efficiently as shown on the spheroid migration trajectories. Fibroblast and MSC spheroids were able to migrate comparable distances (**Fig. 1D**) but MSC clusters moved faster (**Fig. 1E**).

### Mesenchymal cells in cell clusters enable the collective migration of tumor cells

Since CTCCs are generally not composed of only cancer cells, we tested the migration of spheroids composed of a mixture of MCF7 cancer cells with another cell type in a 1:1 ratio (**Fig. 2A**). From now on, we evaluated the efficacy of spheroid migration based on the number of spheroids that migrated to the top of the microwell from their initial position in 24h. Spheroids that moved short distances but did not reach the top of the microwell, were not considered to have migrated by this metric. Cell clusters of MCF7 with MSCs migrated efficiently with MSCs carrying the non-migratory MCF7 cells. Mixed spheroids of cancer cells and fibroblasts showed reduced migratory ability compared to the spheroids composed of fibroblasts alone; nevertheless, migration of these cell clusters was significantly higher than that of mixed cell clusters containing monocytes, macrophages, or ECs (**Fig. 2B,D**).

**Fig. 2.**
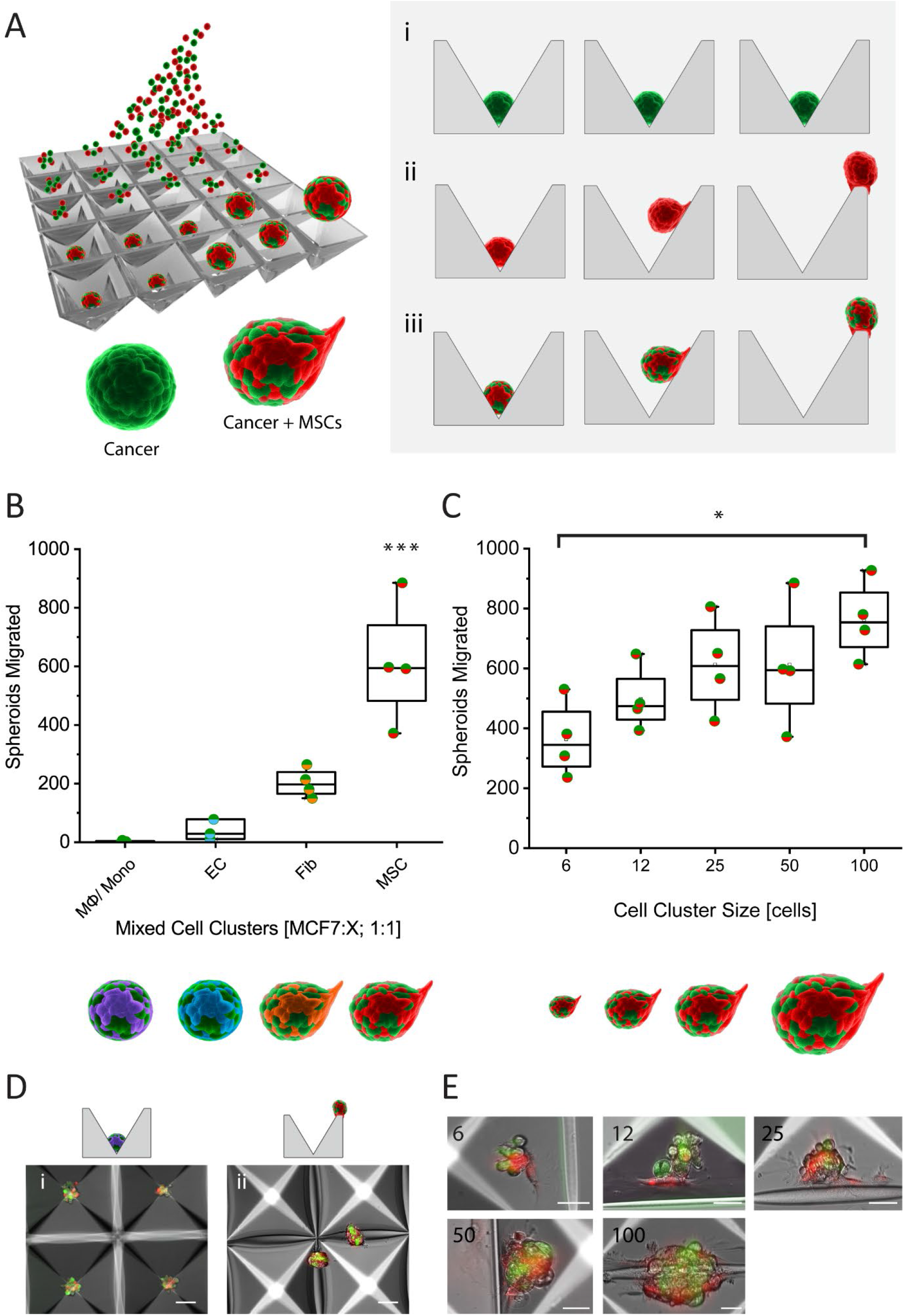
Differences in the migration of mixed spheroids composed of two cell types. **A**. Schematics of spheroid formation in the low adhesive microwells and their migratory behavior. i - non-migratory cancer cell clusters, ii - migratory MSC clusters, iii – mixed spheroids of cancer cells (green) and MSCs (red). **B**. Quantification of mixed spheroid migration. Spheroids in this experiment were formed by MCF7 cancer cells and other cell type in a ratio 1:1. **C**. The migration of mixed cell clusters formed by MCF7 cancer cells and MSCs in the ratio 1:1 was dependent on the size of the cell aggregates. **D**. Visualization of mixed spheroids in the microwells after 24h. i – non-migratory mixed spheroids of macrophages and cancer cells, ii – migratory cell aggregates of MSCs and cancer cells, scale bar – 100 μm. **E**. Cell clusters of 6 to 100 cells formed by cancer cells (green) and MSCs (red) in a ratio 1:1, scale bar – 50 μm.

The size of cell aggregates is another factor that might influence the collective migration. With the increasing size of spheroids composed of a 1:1 ratio of cancer cells and MSCs, more migrating spheroids were detected (**Fig. 2C**), which was apparent when comparing bigger spheroids of 100 cells with cell clusters as small as 6 cells. In all cases, MSCs appeared to carry the cancer cells, which can be seen in **Fig 2E**.

### Migration of tumor cell clusters is enhanced by larger proportions of MSCs in a cluster

To determine if spheroid composition, more specifically, the ratio of cancer cells to MSCs affects cell cluster migratory ability, we prepared mixed cell aggregates that contained 50 cells per spheroid with different ratios of cancer cells to MSCs. Three different cancer cell lines were examined. With increasing counts of MSCs in the spheroid, there was a significant increase in the number of cell clusters capable of migrating to the top of the microwells for all three cancer cell lines (**Fig. 3A,B,C**). Spheroids containing fewer MSCs (10:1 or 5:1 cancer cells:MSCs) were generally the least migratory. Differences among spheroids formed by different cancer cell lines were observed. Cell aggregates with lower numbers of MSCs migrated more when they were mixed with the more aggressive breast cancer cell line MDA-MB-231 compared to spheroids containing more epithelial MCF7 cancer cells.

**Fig. 3.**
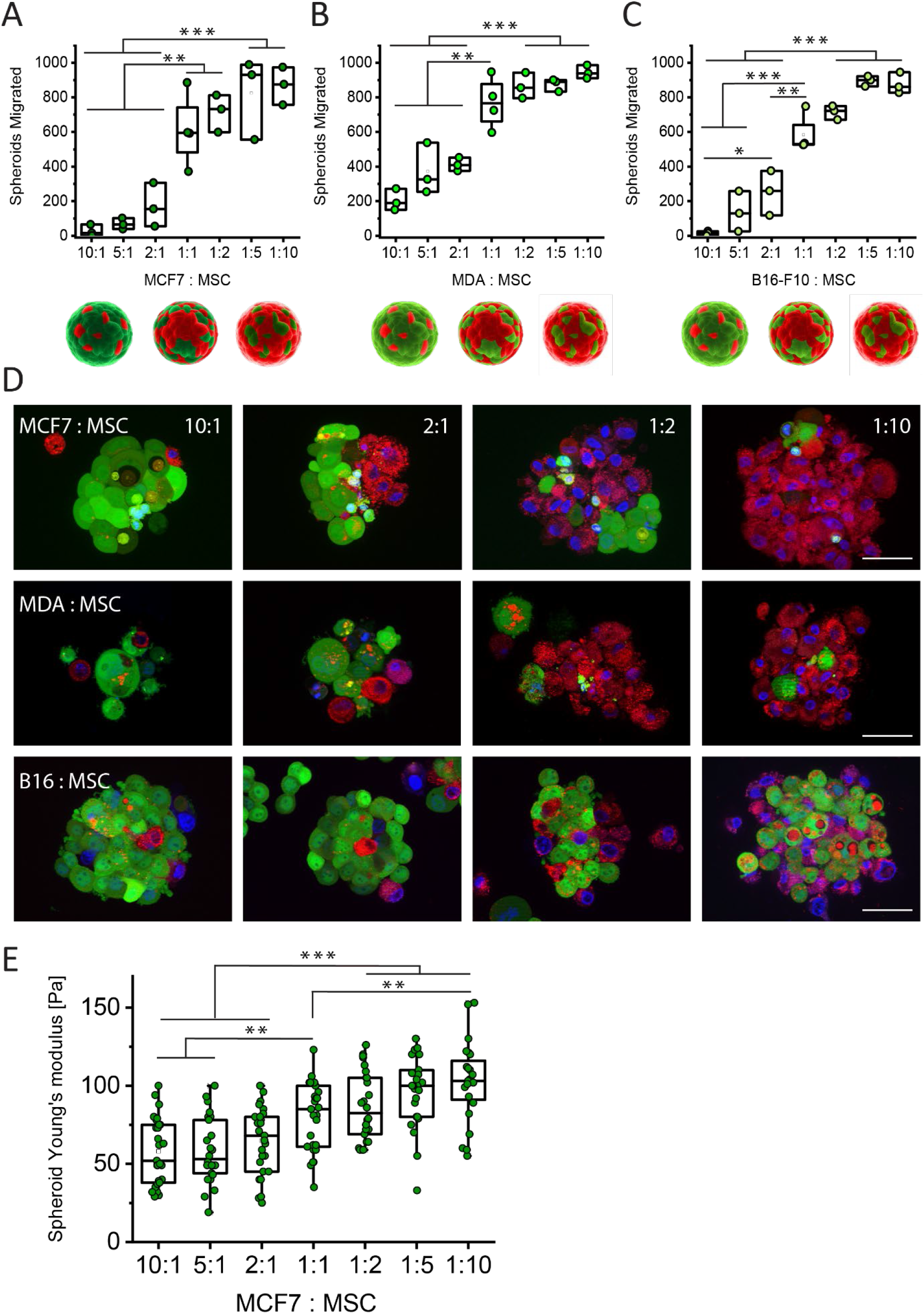
Variation in the cell cluster composition affected cell migration and organization. **A**. Numbers of migrating spheroids composed of different ratios of MCF7 cancer cells and MSCs after 24 hours. **B**. Numbers of spheroids that migrated to the top of the microwell in 24h. Spheroids were composed of different ratios of MDA-MB-231 cancer cells and MSCs. **C**. Numbers of migrating spheroids composed of different ratios of B16-F10 cancer cells to MSCs after 24 hours. **D**. Composition of spheroids formed by MCF7, MDA-MB-231 and B16-F10 cancer cells (green) with MSCs (red) after 2 days in culture. Scale bar – 50 μm. **E**. Differences in Young’s moduli of mixed spheroids composed of different ratios of MCF7 cancer cells to MSCs.

Differences in the compactness and the cell distribution inside the spheroid (**Fig. 3D**) were observed as well. MCF7 cancer cells tend to be more separated from MSCs in the spheroid while B16-F10 cells formed well interconnected structures with MSCs. In contrast to MCF7 cells, monocultures of MDA-MB-231 and B16-F10 cells were not able to form compact spheroids and often migrated in smaller clusters when lower concentrations of MSCs were added. Nevertheless, even in smaller clusters, MSCs were the ones transporting the cancer cells. With increasing numbers of MSCs in the cell aggregates, the spheroids became more compact and migrated as a single entity. In contrast to the conventionally cultured cancer cells in 2D, cancer cells in spheroids showed increased ability to phagocytose MSCs. Internalization of the components derived from MSCs was observed in all cancer cells tested, but prevailed in more aggressive cell lines, mainly in melanoma cancer cells (**Fig. 3D**). In contrast to the other cancer lines tested, the growth kinetics of B16-F10 melanoma cancer cells in the mixed spheroids increased, which altered the ratio of cancer cells to MSCs in favor of melanoma cells.

The changes in spheroid compactness dependent on the ratio of MSCs in the cell cluster can be also documented by differences in the spheroid mechanical properties (**Fig. 3E**). Atomic force microscopy (AFM) revealed that with increasing numbers of MSCs in cell clusters, the spheroids stiffened, which can be related to the differences in the stiffness between cancer cells and MSCs and to the increased compression of the cells due to the forces exerted by MSCs.

### MSC EVs contribute to MSC-mediated migration of tumor cell clusters

In addition to the migratory capability of MSCs, paracrine signals could also play a role in MSC-mediated tumor cell migration. Extracellular vesicles (EVs) are considered to mediate a majority of the paracrine effects of MSCs (28). Therefore, we determined whether EVs secreted by MSCs affected the collective migration of tumor cell clusters. Larger microvesicles were separated from smaller EVs by differential ultracentrifugation, and EVs with a mean size of 70 nm were obtained during the final step of the isolation process after 3-hour ultracentrifugation for 100,000 × g. EVs isolated from ECs, i.e., cells that didn’t support cell aggregate migration in our setup, were used as a control group.

The concentration of EVs secreted by MSCs was on average 10 times higher than the concentration of EVs secreted by ECs (**Fig. 4A**). The vesicular nature of isolated particles was confirmed by transmission electron microscopy (**Fig. 4B**). Isolated EVs were negative for calnexin, a protein localized in endoplasmic reticulum, and positive for EV markers (29) CD63, Flot1, CD81 and CD9 (**Fig. 4C**). The concentration of these markers on EVs varied depending on the cell type.

**Fig. 4.**
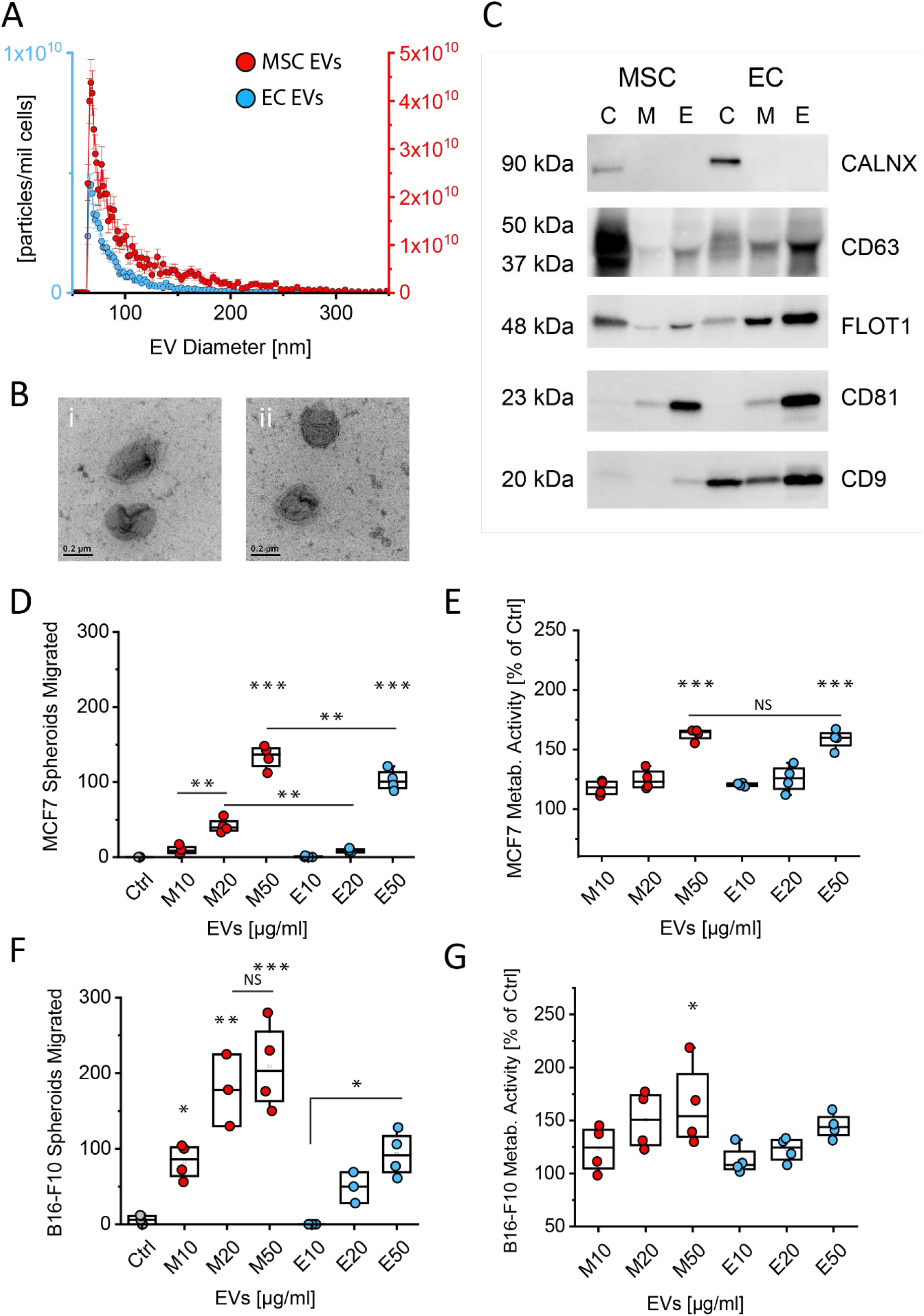
The effect of extracellular vesicles (EVs) secreted by MSCs or ECs on cancer spheroid migration and growth. **A**. EV size and concentration measured by microfluidic resistive pulse sensing. **B**. TEM images of EVs secreted by MSCs (i) and ECs (ii). **C**. Western blotting analysis of EVs secreted by MSCs and ECs, C – cell lysate, M – microvesicles, E – EVs. **D**. Number of migrating MCF7 cancer spheroids after 24-hour incubation with 10, 20 or 50 μg/ml of MSC EVs or EC EVs. Ctrl – control, medium without EVs added. **E**. Metabolic activity of MCF7 spheroids after 48-hour incubation with different concentrations of EVs. **F**. Migration of B16-F10 cancer spheroids after 24-hour incubation with 10, 20 or 50 μg/ml of MSC EVs or EC EVs. Ctrl – control, medium without EVs added. **G**. Metabolic activity of B16-F10 spheroids after 48-hour incubation with different concentrations of EVs.

Cancer aggregates were treated with 3 different concentrations of EVs (10, 20 and 50 μg/ml) and spheroid migration was evaluated after 24 h. Spheroids of two cancer cell lines, MCF7 and B16-F10, were tested. In both cases, we observed the same trend. With increasing concentration of MSC EVs, there was an increase in cancer spheroid migration (**Fig. 4D,F**). Spheroids of more aggressive B16-F10 cancer cells showed higher responsiveness than MCF7 cell clusters to MSC EVs and migrated more even at lower concentrations of MSC EVs. EC EVs were less powerful in promoting cell cluster migration than MSC EVs but an increase in spheroid migration could still be observed, mainly at higher concentrations of EC EVs. Nevertheless, it needs to be pointed out that a million of MSCs produce around 50 μg of EVs in 24 hours but a million of ECs secrete only 10 μg. Therefore, to achieve the same effect as with MSCs, there would need to be significantly higher numbers of ECs present in the tumor. Regardless, the effect of paracrine factors appears to be much weaker compared to the physical presence of MSCs in the co-culture. The migration ability is significantly higher even though the concentration of EVs secreted by MSCs in the spheroids is probably lower than the lowest concentration tested. Both MSC and EC EVs also affected cancer spheroid growth in a dose dependent manner in both cancer spheroids tested (**Fig. 4E,G**). EC EVs increased MCF7 spheroid growth comparably to MSC EVs but in case of B16-F10 spheroids, their effect was less significant in comparison to MSC EVs.

### MSCs facilitated the migration of tumor cell clusters in a 3D microenvironment

To confirm the results obtained from the studies of cell cluster migration in the low-adhesive microwells, we created a micropatterned hydrogel that enabled us to study the spheroid directional collective migration in a 3D microenvironment. For this purpose, cell aggregates were embedded in 5 wt% gelatin methacrylamide (GelMA) hydrogels with aligned stripes of softer and stiffer regions. The hydrogels were polymerized using a visible light photoinitiator, and the stiffness patterns were created by exposing certain regions of the gel to light for different amounts of time using a striped photomask containing 50 μm wide opaque stripes separated by 50 μm wide transparent stripes. Regions receiving longer exposure were stiffer and regions receiving shorter exposure were softer (**Fig. 5A**). In this way, softer, less crosslinked stripes with Young’s modulus ∼ 2kPa and stiffer stripes of 9 kPa were created (**Fig. 5C**). In this system, the migration of cell aggregates composed of MCF7 cancer cells was compared with the behavior of cancer cell clusters treated with MSC EVs at the concentration of 50 μg/ml and with mixed spheroids of MCF7 cells with MSCs in the ratio 1:1. The difference in spheroid area and circularity was evaluated after 7 days of culture. The growth of cancer cell aggregates in the microenvironment containing MSC EVs was slightly increased compared to the cancer spheroids alone. The largest spheroid area was observed in the mixed cell aggregates when MSCs were physically present in the spheroids (**Fig. 5E**). Cell cluster directional migration was evaluated based on the change in circularity from the initial spheroids. Cancer cell aggregates or spheroids treated with MSC EVs did not show significant change in circularity, however, mixed spheroids containing MSCs elongated over the course of 7 days as they migrated in the direction of stripes (**Fig. 5F**). Interestingly, all aggregates in this 3D microenvironment migrated only in the softer, less crosslinked regions with MSCs functioning as a tip cells, guiding the migration of the whole cluster. The rest of the MSCs were also observed to cover the cancer cells at the interface between the cells and the microenvironment (**Fig. 5Dii**).

**Fig. 5.**
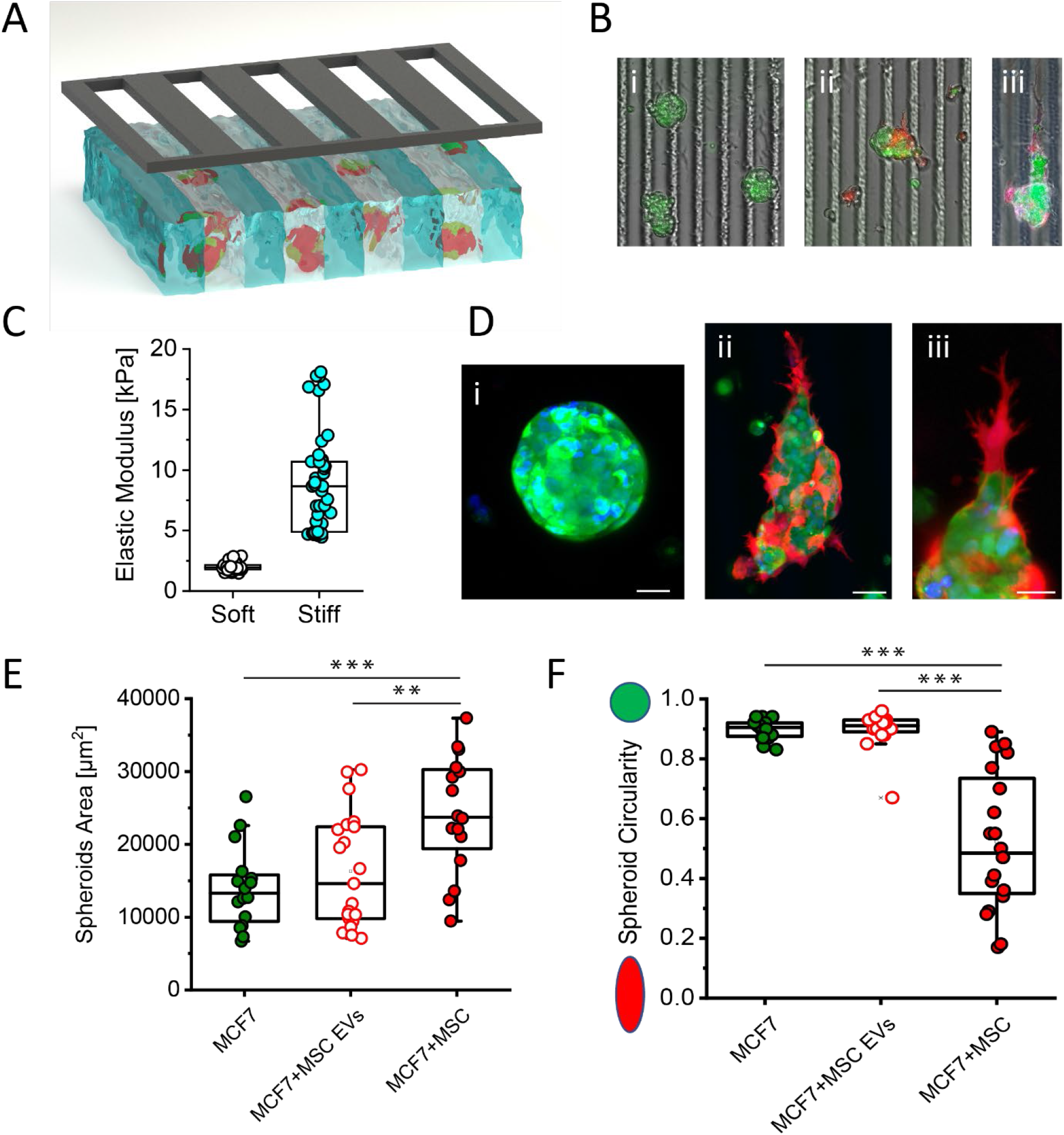
Cell cluster behavior in the micropatterned hydrogel with stripes of two different stiffnesses. **A**. Schematics of micropatterned GelMA hydrogel with embedded spheroids fabricated with the use of photomask that blocks UV irradiation of softer stripes. **B**. Visualization of spheroids embedded in the striped hydrogel. i – MCF7 cancer spheroids after 7 days in culture, ii – MCF7+MSC spheroid after 2 days, and iii - MCF7+MSC spheroid after 7 days in culture, with MSCs acting like tip cells guiding the migration. **C**. Elastic modulus of stripes of different stiffnesses measured by AFM. **D**. Confocal images of spheroids after 7 days in culture. i – MCF7 spheroid, ii – MCF7+MSC spheroid, scale bar – 50 μm, iii – detail of MSC tip cell in MCF7+MSC spheroid, scale bar – 25 μm. **E**. Area of spheroids after 7 days in culture. **F**. Circularity of spheroids after 7 days in culture suggesting directional movement of MCF7+MSC mixed spheroids.

Together, these results indicate that MSCs might participate in the dissemination of less migratory cancer cells (e.g., cancer stem cells) in vivo by transporting cell clusters and secreting paracrine factors, which are less potent in inducing the migration but might support the cancer cell survival and proliferation.

## Discussion

Recent reports on the increased metastatic potential of CTCCs compared to single cancer cells has highlighted the importance of studying the process of collective cell migration. However, this process, mainly in the context of cancer, is generally very slow (10 to 15 μm per day (30)) and the conditions that promote CTCC migration are not well understood. In vivo tracking of migrating cell clusters is very challenging. Therefore, *in vitro* systems that enable the study of collective cell migration are of a great interest.

Factors such as substrate stiffness, substrate geometry, and cell adhesion ligand concentration are known to influence migration of single cells *in vitro*. Cells tend to elongate and migrate along topographical features such as microgrooves or fibrillar structures (31). Moreover, curvature dependent changes in cell behavior were also observed on surfaces containing convex and concave features, where cells in the concave regions showed decreased cell area and faster migration compared to cells on flat or convex structures, which has been attributed to increased cytoskeletal contractility and changes in nuclear geometry (32) (33). In addition to substrate geometry, cell-substrate adhesion regulates dynamic cell motility. Usually, low levels of adhesion do not support cell migration, while moderate adhesion promotes dynamic cell motility and high adhesion suppresses cell locomotion since cells form more bonds with the substrate, which hinder their detachment from the substrate (34, 35).

In this study, we examine cell aggregate migration in low-adhesion inverted-pyramidal microwells. Spheroids seeded on a flat low-adhesive surface that was treated with the same anti-adhesive solution as the microwells did not show as significant migratory properties as cell clusters on patterned surfaces. Spheroid formation and migration in the microwells are processes connected with increased EV secretion (which can be best observed on the video of large cell aggregate migration (**Supplemental video S3**), which might contribute to the establishment of a gradient in the microwell that then affects spheroid directional movement.

So far, there are just few articles focused on cell aggregate migration, where cell cluster migration was observed either on soft 2D surfaces (36) or in soft 3D gels (37). However, spheroid migration in both of these systems was slow and the clusters needed to be followed by time-lapse microscopy, which does not allow for high-throughput screening of many cell aggregates at once. On the contrary, our system permits us to evaluate migration simply by the end-point assessment of the spheroid migratory ability based on the fact that the cell cluster reached the top of the microwell and stayed at this stationary state.

Migration of aggregates composed of various cell types tested with the use of this system showed striking differences in their motility. Leukocyte migration is generally considered adhesion-independent. However, neither monocytes, nor macrophages migrated on these micropatterned surfaces. This might be due to the lack of confinement that is thought to be necessary for the efficient amoeboid migration (38). We did not observe any significant motility of cancer, epithelial or endothelial spheroids. Clusters containing mesenchymal cells migrated efficiently as a majority of the clusters reached the top edge of the microwell in less than 24 hours.

Cells have been reported to be able to transport cargos such as cellular backpacks, which are engineered particles conjugated to the cell surface (39). However, there are no reports on the “cell taxi service” that can transport other cells. While the phenomenon of mesenchymal stem cells transporting cancer cells was visualized *in vitro*, it suggests that the cancer spreading might not be a process solely dependent just on the migratory abilities of cancer cells. This is also supported by the observation that the administration of selected subpopulation of highly migratory cancer cells did not promote more aggressive tumor growth or increased metastases (40).

We have also observed the dependence of the motility on the size of the cluster, where larger cell aggregates migrated more efficiently than smaller ones. This is in agreement with other studies, such as clusters of T cells which were observed to move more slowly but with increased directionality comparing to single cells (41).

Besides the cluster size, cell aggregate composition influences its migratory ability. Highlighted in this study is the ability of MSCs to make the spheroid more compact so that even the otherwise loosely adhesive cancer cells such as B16-F10 or MDA-MB-231 are migrating in collectives and not as single cells. Moreover, it was observed that in contrast to 2D cultures, cancer cells in 3D spheroids without contact to the surrounding ECM readily internalize components derived from MSCs or even engulf entire cells, which can give them further survival advantage in the form of nutrients derived from MSCs (42).

EVs secreted by MSCs enhanced cancer spheroid metabolic activity by approx. 60% and increased the number of migrating cancer aggregates by 13-20% at the highest concentrations. The percentage of migrating cancer spheroids treated by MSC EVs is significantly lower in comparison to the 60% increase in numbers of migrating spheroids in the co-culture model. It thus seems that the paracrine factors, such as EVs, contribute in part to the overall pro-migratory effect of MSCs but that the physical presence of MSCs in the cell cluster has a higher impact. However, the effect of stromal EVs might be further multiplied *in vivo.* For example, EV internalization by immune cells might induce additional reactions that cannot be addressed by the two-component *in vitro* system.

We were further interested if the behavior observed in the 2.5D environment on the micropatterned surfaces reflects what is happening in the 3D microenvironment. For this purpose, micropatterned 3D gels were designed with micropatterned crosslink density. An abundance of research has shown that the stiffness of the cellular microenvironment influences the migratory properties of cancer cells. However, these models lack the *in vivo* complexity of the real tumor microenvironment that has quite heterogenous stiffness (43), which might affect the cancer cell migration. Our *in vitro* system does not attempt to model the entire complexity of the tumor microenvironment but it is based on the observation that the collective cell migration is induced in confined spaces (44). By controlling the crosslinking of the GelMA gel, we created stripes with more and less crosslinking and we observed a higher collective migration in the softer regions. However, this was not solely the property of cancer spheroids but the same behavior was observed also for other cell types, such as mixed spheroids composed of endothelial cells and MSCs (results not shown). This observation is in agreement with a previously published study showing that cells choose a path of lowest energetic cost while navigating through a heterogenous microenvironment (45). In the co-cultures with MCF7 cancer cells, MSCs were always present at the leading edge of the migrating cell cluster, while the rest of MSCs surrounded the cancer cells in the core of the spheroid.

The collective migration guided by stripes of different stiffnesses highlights the importance to produce the morphological asymmetry in the microenvironment, e.g. by the confinement, gradient or other means that might enable to study the collective migration of cell clusters in 3D. When spheroids are embedded in a gel of uniform stiffness, no directional migration could be observed. Thus, this study emphasizes the importance of symmetry breaking by different means to study the collective migration *in vitro* and advance our understanding of this interesting phenomenon that is important not only for the cancer spreading but also in the development.

## Conclusion

The generally accepted theory of cancer cell spreading is based on the notion that cancer cells need to change their phenotype by epithelial-to-mesenchymal transition and acquire mesenchymal features in order to become more migratory and invasive. In this study, our findings suggest that MSCs or other mesenchymal cells such as fibroblasts may serve as carriers to transport the low-migratory tumor cells. In addition, MSCs secrete paracrine factors (such as EVs) to promote cancer cell growth and invasion. This study highlights the need to better understand the process of collective cell migration and the contribution of the tumor stroma to the cancer progression. Furthermore, it also implies the risks of using MSCs in regenerative therapies after tumor removal as it might lead to the increased metastatic spreading of cancer cells.

## Materials and Methods

### Chemicals and biologicals

Unless noted otherwise, all chemicals were purchased from Sigma-Aldrich, Inc. Cell culture reagents, solutions, and dishes were obtained from Thermo Fisher Scientific, except as indicated otherwise. Gelatin methacrylamide was fabricated using gelatin from fish (J. T. Baker, 250 bloom) as previously described (46).

### Cells

Cancer cell lines MCF-7 and MDA-MB-231 are from the American Type Culture Collection (ATCC, Manassas, VA). GFP expressing B16-F10 cells were obtained from Creative Biogene (Shirley, NY, USA). Unless otherwise stated, all cells were cultured in Dulbecco’s Modified Eagle’s Medium (DMEM) with 10% FBS (Gibco) and 1% penicillin/streptomycin (Gibco). RAW 264.7 macrophages as well as THP-1 monocytes were obtained from ATCC (Manassas, VA). THP-1 were cultured in RPMI-1640 medium with 10% FBS. Primary human dermal fibroblasts as well as human umbilical vein endothelial cells (ECs) were purchase from ATCC. Endothelial cells were cultured in EGM-2 medium (Lonza). Human mesenchymal stem cells (hMSCs) were obtained from Texas A&M Health Science Centre College of Medicine. MSCs and ECs in a passage 3-5 were used in this study. In short-term co-cultures, MSCs were stained with red and cancer cells with green CellTracker (Invitrogen) according to the manufacturer’s instructions. For long-term co-cultures, MSCs were transduced to stably express LifeAct-RFP using the lentiviral vector (pTwist Lenti CMV Puro) backbone (Twist Bioscience). MCF7 cancer cells were transduced to stably express LifeAct-GFP using the same lentiviral vector backbone.

### Spheroid formation and migration

The majority of the microwell experiments were performed using the AggreWell 400 (STEMCELL Technologies) microwell dishes. Microwells were incubated with Anti-Adherence Rinsing Solution (STEMCELL Technologies) for 5 min, followed by 5 min centrifugation at 2000 x g. The wells were washed twice with medium without FBS then filled with 0.5ml complete medium containing 10% FBS. To form spheroids containing 50 cells each, 60 000 cells were seeded per well for a 24 well plate with a final volume of 1ml of medium containing 10% FBS. Seeded cells were centrifuged for 2min at 100 x g and an even cell distribution at the bottom of the microwells was confirmed under the microscope. To evaluate the migration of spheroids formed by different cell types, time-lapse microscopy was performed at a rate of one frame every 4min. Spheroid migration was manually tracked using ImageJ and evaluated using the Chemotaxis and Migration Tool (Ibidi). In other experiments, spheroid migration was evaluated by counting the number of spheroids that reached the top of the microwell after 24h. Cell metabolic activity was measured using the PrestoBlue Reagent (Invitrogen) according to the manufacturer’s instructions.

### AFM of spheroids

For mechanical characterization of the spheroids, standard V-shaped gold-coated silicon nitride tipless AFM cantilevers (BrukerNano, Camarillo, CA) were used after modification with latex beads (10 μm diameter). The cantilever spring constant was measured using the thermal fluctuations method. The spring constant of the cantilevers used in this work was found to be 0.134 N/m. Measurements were performed using a Veeco AFM II Dimension 3100 (Veeco Metrology Inc., now BrukerNano) instrument in liquid mode. The force displacement curves were recorded with a vertical ramp size of 5 μm. To minimize viscoelastic effects, force-indentation curves were recorded at a frequency of 1 Hz.

### EV isolation

EV-depleted FBS was obtained by 18h ultracentrifugation at 100, 000 x g, 4 °C. Cells were washed with PBS and cultured in medium containing EV-depleted FS for 24h. For all EV isolations, cell viability was higher than 95%. Conditioned medium containing EVs was centrifuged at 2, 000 x g for 10min, filtered through 0.22 μm filter, concentrated with a 10kDa Centricon Plus-70 centrifugal filter (UFC701008, Sigma Millipore) and centrifuged at 100,000 x g for 3 h at 4 °C. The EV pellet was resuspended in 100 μl PBS and stored in −80 °C until further use.

#### EV Characterization

### Western blot

Cells were lysed in RIPA buffer with protease inhibitors (HALT™ Protease Inhibitor Cocktail, EDTA-free (100X), Thermo Scientific, 87785) for 20 min on ice, and were then centrifuged for 15 min at 14,000 g. The protein concentration was measured by Micro-BCA (Thermo Scientific, 23235) in the presence of 0.2 % SDS. For SDS-PAGE, samples were mixed with Laemmli Sample Buffer, boiled for 5 min at 95°C, and separated on 4-20% gradient polyacrylamide gels (4561094, Bio-Rad) before they were transferred to the PVDF membranes. The membranes were blocked with 5% non-fat milk for 1h and were then incubated overnight at 4°C with primary antibodies: anti-CANX (1:500, rabbit polyclonal anti-human ABclonal, A15631); anti-CD63 (1:500, mouse monoclonal anti-human CD63 antibody, MEM-259, abcam, ab8219); anti-FLOT1 (1:500, rabbit polyclonal anti-human ABclonal, A6220); anti-CD81 (1:300, mouse monoclonal anti-human CD81 antibody, Invitrogen, MA5-13548); anti-CD9 (1:500, mouse monoclonal anti-human CD9, clone MM2/57, Invitrogen, AHS0902). The membranes were then washed three times with 0.1% Tween 20 in PBS for 5 minutes at RT, and were incubated with secondary antibody conjugated to horseradish peroxidase (donkey anti-rabbit IgG-HRP, sc-2313, or donkey anti-mouse IgG-HRP, sc-2314, Santa Cruz Biotechnology) for 2h at 4°C. After the membranes had been washed three times with 0.1% Tween 20 in PBS, proteins were visualized with a chemiluminescence substrate (SuperSignal West Femto Maximum Sensitivity Substrate, Thermo Scientific, 34095).

### Transmission electron microscopy (TEM)

Undiluted suspensions of EVs were suspended on grids with a thin formvar/carbon film and were allowed to adsorb for 20 minutes. Excess liquid was blotted away with filter paper and the grids were then washed three times with 20mM HEPES and 150mM NaCl and were negatively stained with 0.4% uranyl acetate, 3% methylcelulose for 1 minute. Excess solution was blotted away, and the grids were air-dried. Samples were imaged using JEOL JEM 1200 EX operated at 120 kV.

### Microfluidic Resistive Pulse Sensing

The concentration and the size distribution of EVs was analyzed by microfluidic resistive pulse sensing using Spectradyne nCS1 (Spectradyne LLC) with C-400/TS-400 cartridge.

### Fabrication and characterization of hydrogels with micropatterned stiffness

Hydrogels were covalently adhered to a glass substrate as previously described (47). Briefly, round pieces of coverglass (12 mm diameter, Fisher) were functionalized with methacrylate groups using the following procedure. The coverglass was first cleaned activated with oxygen plasma (Plasma Prep II, SPI Supplies) for 5 minutes. The activated coverglass was then reacted with 3-(trimethoxysilyl)propyl methacrylate (0.5 mL) in 25 mL of an ethanol and acetic acid solution for 30 minutes. The coated coverglass was then washed with methanol three times before drying. Silanized coverglass pieces were used within two hours of functionalization.

Glass slides were rendered hydrophobic, and thus non-adherent to the polymerized hydrogels, by wiping a drop of Gel Slick (Lonza) on the slide surface. This process was repeated once to ensure a sufficient coating. The slides were then briefly rinsed with DI water to clean the surface and dried before use.

Photopolymerizable gelatin-based hydrogels were prepared using gelatin methacrylamide (GelMA) and a visible-light photoinitiator lithium phenyl-2,4,6-trimethylbenzoylphosphinate (LAP). GelMA stock solutions were heated in a 55° C water bath until dissolved and then centrifuged to remove any bubbles. Since the gelatin used was fish-based, once dissolved the stock solutions remained liquid at room temperature for approximately one hour. If the gelatin stock solutions underwent physical gelation, they could easily be re-liquefied in a 40° water bath. A pre-polymer solution of 5.0 w/w% GelMA with 0.10 w/w% LAP in PBS was prepared and pipetted (10 μL/gel) between a hydrophobically modified glass slide and 12 mm round glass coverslips functionalized with methacrylate groups that were separated by 200 μm spacers. For spheroid encapsulation studies, spheroids were mixed with the pre-polymer solutions, while maintaining 5.0 w/w% GelMA and 0.10 w/w% LAP and immediately polymerized with spheroids in situ. The gels were polymerized, and thus chemically crosslinked, into a hydrogel using a collimated light source (EXFO Omnicure S1000) filtered with a 405 nm bandpass filter (Newport) with an output intensity of 4.8 mW/cm^2^ as measured by a spectroradiometer (International Light Technologies, ILT950) between 350 and 500 nm. Photomasks (CAD/Art Services) with stripes of 50 μm opaque regions and 50 μm wide transparent regions were used to block light from reaching certain regions of the gels so that certain sections received 37.5 seconds exposure while others received 150 seconds exposure. These spatially defined differences in photopolymerization time led differences in crosslink density, thus, gels with spatially defined differences in elastic modulus were produced. After polymerization, the coverglass (with the attached hydrogel) was pried off the glass slide, and submersed in cell culture media. Collective migration of cells in the micropatterned gels was evaluated after 7 days when spheroids were fixed with 4% paraformaldehyde and permeabilized with 0.05% Triton X-100 and 1% bovine serum albumin solution for 15 min at room temperature. Nuclei were stained with DAPI. Images were taken on Leica SP8 confocal microscope and were evaluated using ImageJ.

To determine the mechanical properties of the hydrogels with micropatterned stiffness, hydrogels without spheroids were prepared as described above and submersed in 1× PBS to swell overnight. The elastic modulus of both 150 second (stiff) and 37.5 second (soft) exposed regions were measured using Atomic Force Microscopy (AFM) in PBS using a JPK Nanowizard 4a BioScience AFM using the force spectroscopy mode in the Nano and Pico Characterization Laboratory at California Nanosystems Institute, UCLA. A CP-qp-CONT-SiO-B probe with a 3.5 μm diameter SiO2 sphere (sQube®) was used to indent the samples. For the quantitative measurements of Young’s modulus, the spring constant of the cantilever was measured using the AFM’s internal contact-free thermal tuning method. Single indentations were performed with a total force of 4.0 nN. At least 15 indentations were performed at 15 different locations across the gel surface for each region. The photomasks contained fiduciary markers so that the softer and stiffer regions of the hydrogels could be correctly identified. All AFM force curve analysis was performed using the JPK Data Processing software. The Young’s modulus was calculated by using a Hertz/Sneddon spherical fit with a Poisson’s ratio of ν = 0.5 (48).

## Statistical analysis

One-way ANOVA with Tukey’s post-hoc test was used to determine significant differences across multiple samples (*** p < 0.001, ** p < 0.01, * p < 0.05).

## Acknowledgments

This work was supported in part by grants from the National Institute of Health (R56DE029157), the California Institute for Regenerative Medicine (Grant Number DISC2COVID19-11838), and Eli and Edythe Broad Center of Regenerative Medicine and Stem Cell Research Award Program. S.C.P.N gratefully acknowledges support from a Ruth L. Kirschstein Predoctoral Fellowship (NIH-F31DE026356).

---

**Suppl. Fig. 1.**
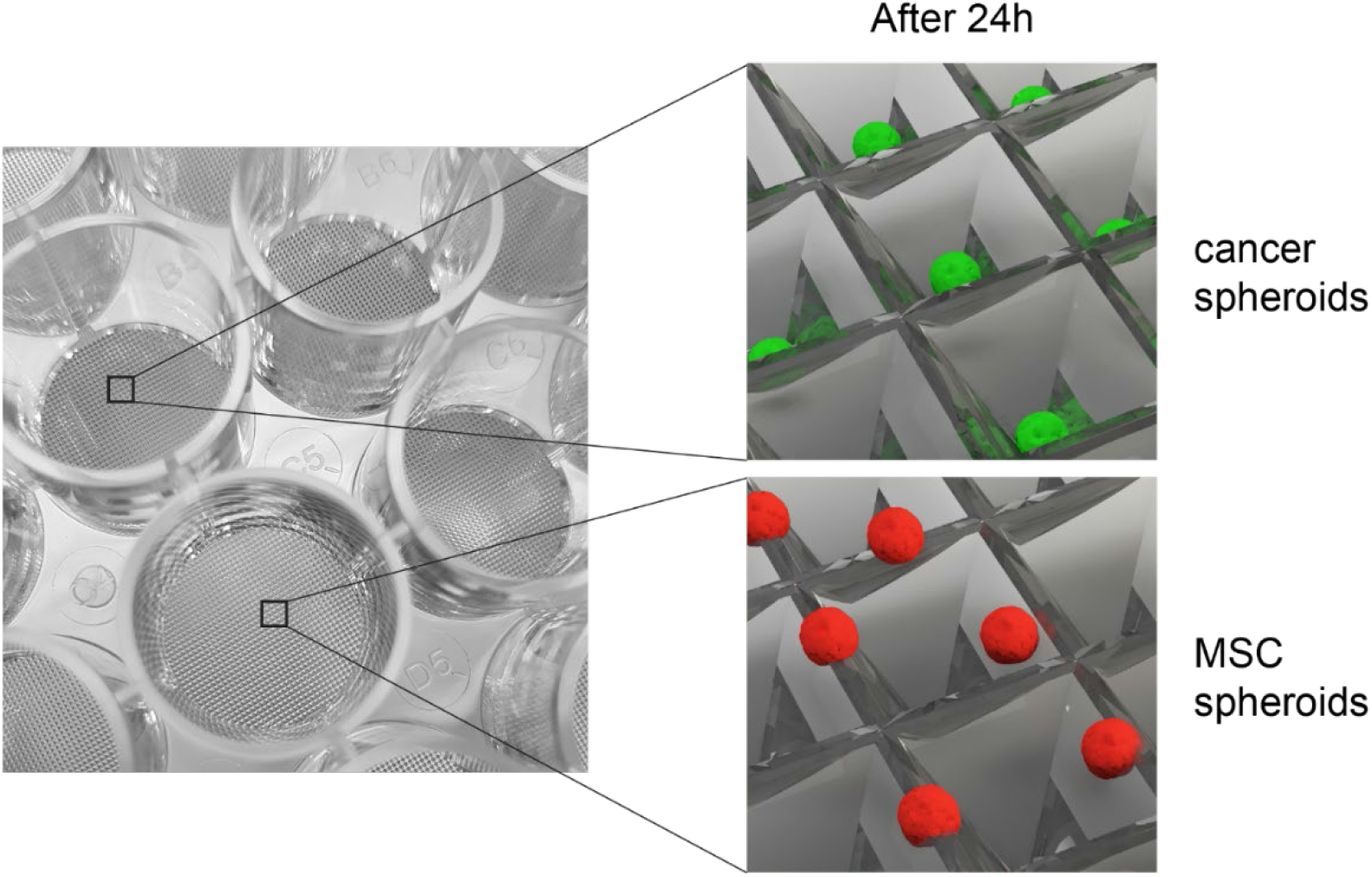
Visualization of differences in cancer and MSC spheroid migration after 24h in inverted-pyramidal microwells

## References

1. Dillekås H, Rogers MS, & Straume O (2019) Are 90% of deaths from cancer caused by metastases? Cancer Med 8(12):5574–5576.

2. Valastyan S & Weinberg RA (2011) Tumor metastasis: molecular insights and evolving paradigms. Cell 147(2):275–292.

3. Cheung KJ & Ewald AJ (2016) A collective route to metastasis: Seeding by tumor cell clusters. Science 352(6282):167–169.

4. Cheung KJ, et al. (2016) Polyclonal breast cancer metastases arise from collective dissemination of keratin 14-expressing tumor cell clusters. Proceedings of the National Academy of Sciences 113(7):E854–E863.

5. Aceto N, et al. (2014) Circulating tumor cell clusters are oligoclonal precursors of breast cancer metastasis. Cell 158(5):1110–1122.

6. Maddipati R & Stanger BZ (2015) Pancreatic Cancer Metastases Harbor Evidence of Polyclonality. Cancer Discovery 5(10):1086–1097.

7. Au SH, et al. (2016) Clusters of circulating tumor cells traverse capillary-sized vessels. Proceedings of the National Academy of Sciences of the United States of America 113(18):4947–4952.

8. Hong Y, Fang F, & Zhang Q (2016) Circulating tumor cell clusters: What we know and what we expect (Review). Int J Oncol 49(6):2206–2216.

9. Duda DG, et al. (2010) Malignant cells facilitate lung metastasis by bringing their own soil. Proceedings of the National Academy of Sciences 107(50):21677–21682.

10. Cuiffo BG & Karnoub AE (2012) Mesenchymal stem cells in tumor development. Cell Adhesion & Migration 6(3):220–230.

11. Spaeth E, Klopp A, Dembinski J, Andreeff M, & Marini F (2008) Inflammation and tumor microenvironments: defining the migratory itinerary of mesenchymal stem cells. Gene Therapy 15(10):730–738.

12. Chaturvedi P, et al. (2013) Hypoxia-inducible factor–dependent breast cancer–mesenchymal stem cell bidirectional signaling promotes metastasis. The Journal of Clinical Investigation 123(1):189–205.

13. Krueger TEG, Thorek DLJ, Denmeade SR, Isaacs JT, & Brennen WN (2018) Concise Review: Mesenchymal Stem Cell-Based Drug Delivery: The Good, the Bad, the Ugly, and the Promise. STEM CELLS Translational Medicine 7(9):651–663.

14. Lee H-Y & Hong I-S (2017) Double-edged sword of mesenchymal stem cells: Cancer-promoting versus therapeutic potential. Cancer Sci 108(10):1939–1946.

15. Roccaro AM, et al. (2013) BM mesenchymal stromal cell–derived exosomes facilitate multiple myeloma progression. The Journal of Clinical Investigation 123(4):1542–1555.

16. Zhu W, et al. (2012) Exosomes derived from human bone marrow mesenchymal stem cells promote tumor growth in vivo. Cancer Letters 315(1):28–37.

17. Zhang T, et al. (2013) Bone marrow-derived mesenchymal stem cells promote growth and angiogenesis of breast and prostate tumors. Stem Cell Research & Therapy 4(3):70.

18. Patel SA, et al. (2010) Mesenchymal Stem Cells Protect Breast Cancer Cells through Regulatory T Cells: Role of Mesenchymal Stem Cell-Derived TGF-β. The Journal of Immunology 184(10):5885–5894.

19. Biswas S, et al. (2019) Exosomes Produced by Mesenchymal Stem Cells Drive Differentiation of Myeloid Cells into Immunosuppressive M2-Polarized Macrophages in Breast Cancer. J Immunol 203(12):3447–3460.

20. Shinagawa K, et al. (2010) Mesenchymal stem cells enhance growth and metastasis of colon cancer. International Journal of Cancer 127(10):2323–2333.

21. Kalluri R (2016) The biology and function of fibroblasts in cancer. Nature Reviews Cancer 16(9):582–598.

22. Karnoub AE, et al. (2007) Mesenchymal stem cells within tumour stroma promote breast cancer metastasis. Nature 449(7162):557–563.

23. Zhong W, et al. (2017) Mesenchymal stem cells in inflammatory microenvironment potently promote metastatic growth of cholangiocarcinoma via activating Akt/NF-κB signaling by paracrine CCL5. Oncotarget 8(43):73693–73704.

24. Ridge SM, Sullivan FJ, & Glynn SA (2017) Mesenchymal stem cells: key players in cancer progression. Molecular Cancer 16(1):31.

25. Stevens K, et al. (2013) InVERT molding for scalable control of tissue microarchitecture. Nature communications 4(1):1–11.

26. Nguyen AH, Wang Y, White DE, Platt MO, & McDevitt TC (2016) MMP-mediated mesenchymal morphogenesis of pluripotent stem cell aggregates stimulated by gelatin methacrylate microparticle incorporation. Biomaterials 76:66–75.

27. Mosaad E, Chambers K, Futrega K, Clements J, & Doran M (2018) The Microwell-mesh: A high-throughput 3D prostate cancer spheroid and drug-testing platform. Scientific reports 8(1):1–12.

28. Witwer KW, et al. (2019) Defining mesenchymal stromal cell (MSC)-derived small extracellular vesicles for therapeutic applications. Journal of extracellular vesicles 8(1):1609206–1609206.

29. Thery C, et al. (2018) Minimal information for studies of extracellular vesicles 2018 (MISEV2018): a position statement of the International Society for Extracellular Vesicles and update of the MISEV2014 guidelines. J Extracell Vesicles 7(1):1535750.

30. Weigelin B, Bakker G-J, & Friedl P (2012) Intravital third harmonic generation microscopy of collective melanoma cell invasion. IntraVital 1(1):32–43.

31. Hennig K, et al. (2020) Stick-slip dynamics of cell adhesion triggers spontaneous symmetry breaking and directional migration of mesenchymal cells on one-dimensional lines. Science Advances 6(1):eaau5670.

32. Werner M, et al. (2017) Surface Curvature Differentially Regulates Stem Cell Migration and Differentiation via Altered Attachment Morphology and Nuclear Deformation. Advanced Science 4(2):1600347.

33. Park JY, Lee DH, Lee EJ, & Lee S-H (2009) Study of cellular behaviors on concave and convex microstructures fabricated from elastic PDMS membranes. Lab on a Chip 9(14):2043–2049.

34. Palecek SP, Loftus JC, Ginsberg MH, Lauffenburger DA, & Horwitz AF (1997) Integrin-ligand binding properties govern cell migration speed through cell-substratum adhesiveness. Nature 385(6616):537–540.

35. Cui K, Ardell CL, Podolnikova NP, & Yakubenko VP (2018) Distinct Migratory Properties of M1, M2, and Resident Macrophages Are Regulated by α(D)β(2) and α(M)β(2) Integrin-Mediated Adhesion. Frontiers in immunology 9:2650–2650.

36. Beaune G, et al. (2018) Spontaneous migration of cellular aggregates from giant keratocytes to running spheroids. Proceedings of the National Academy of Sciences 115(51):12926–12931.

37. Zajac O, et al. (2018) Tumour spheres with inverted polarity drive the formation of peritoneal metastases in patients with hypermethylated colorectal carcinomas. Nature Cell Biology 20(3):296–306.

38. Liu Y-J, et al. (2015) Confinement and Low Adhesion Induce Fast Amoeboid Migration of Slow Mesenchymal Cells. Cell 160(4):659–672.

39. Shields CW, et al. (2020) Cellular backpacks for macrophage immunotherapy. Science Advances 6(18):eaaz6579.

40. Amaro A, et al. (2016) A highly invasive subpopulation of MDA-MB-231 breast cancer cells shows accelerated growth, differential chemoresistance, features of apocrine tumors and reduced tumorigenicity in vivo. Oncotarget 7(42):68803–68820.

41. Malet-Engra G, et al. (2015) Collective Cell Motility Promotes Chemotactic Prowess and Resistance to Chemorepulsion. Current Biology 25(2):242–250.

42. Overholtzer M & Brugge JS (2008) The cell biology of cell-in-cell structures. Nature Reviews Molecular Cell Biology 9(10):796–809.

43. Plodinec M, et al. (2012) The nanomechanical signature of breast cancer. Nature Nanotechnology 7(11):757–765.

44. Haeger A, Wolf K, Zegers MM, & Friedl P (2015) Collective cell migration: guidance principles and hierarchies. Trends in Cell Biology 25(9):556–566.

45. Zanotelli MR, et al. (2019) Energetic costs regulated by cell mechanics and confinement are predictive of migration path during decision-making. Nature Communications 10(1):4185.

46. Norris SCP, Delgado SM, & Kasko AM (2019) Mechanically robust photodegradable gelatin hydrogels for 3D cell culture and in situ mechanical modification. Polymer Chemistry 10(23):3180–3193.

47. Norris SCP, Soto J, Kasko AM, & Li S (2021) Photodegradable Polyacrylamide Gels for Dynamic Control of Cell Functions. ACS Appl Mater Interfaces.

48. Mott PH & Roland CM (2009) Limits to Poisson’s ratio in isotropic materials. Physical Review B 80(13):132104.

